## Supplementary figs and tables for "High-resolution micro-epidemiology of parasite spatial and temporal dynamics in a high malaria transmission setting in Kenya"

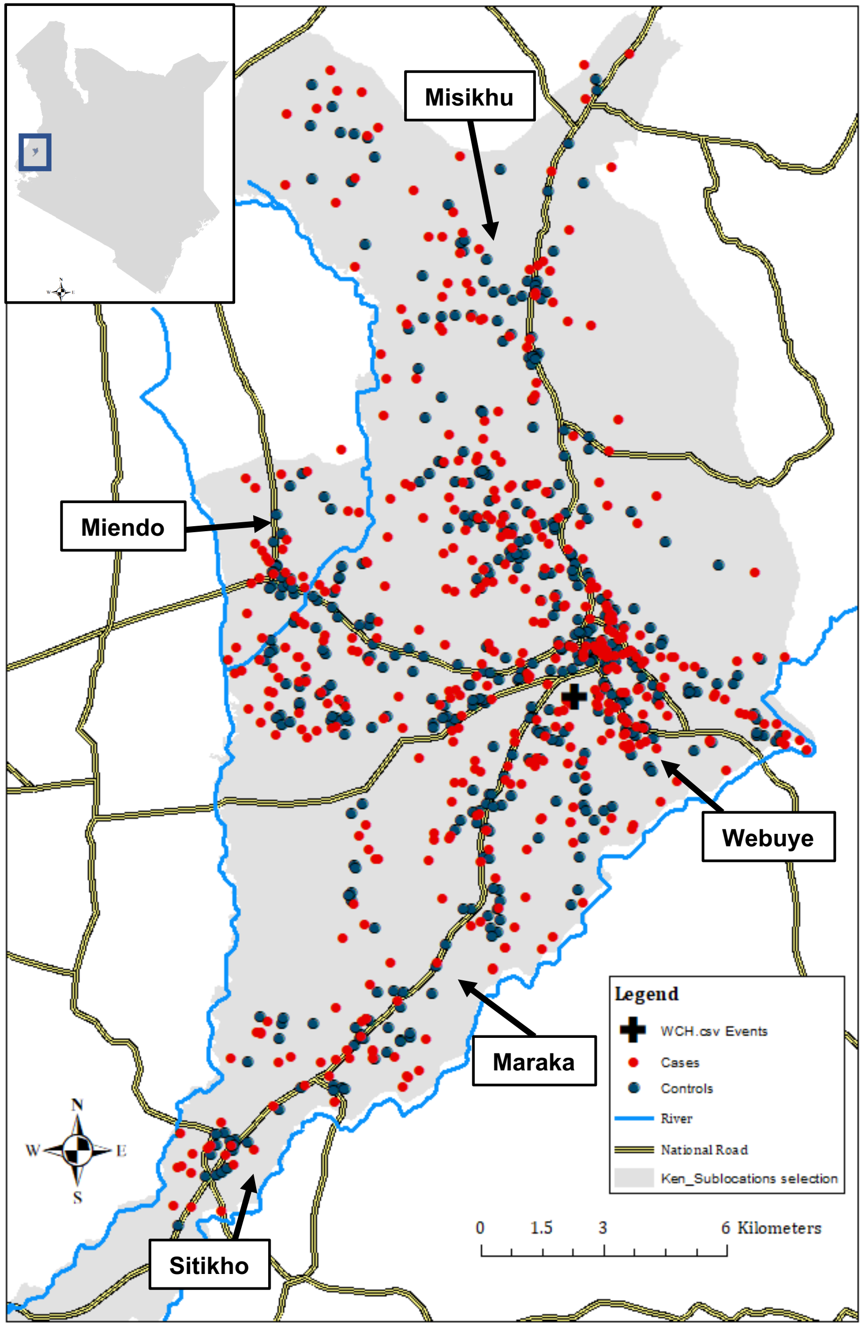

**Supplementary Fig. 1** Map of study area in Bungoma county, western Kenya. Black cross indicates Webuye County Hospital (WCH), and gray outline denotes 10 administrative locations adjacent to the hospital. The 5 locations with the highest case incidence are labelled. The geographic location of the households of case and control children are indicated as red and blue dots, respectively. Blue denotes rivers, and yellow national roads. Visualization created using ArcGIS, version 10.7.

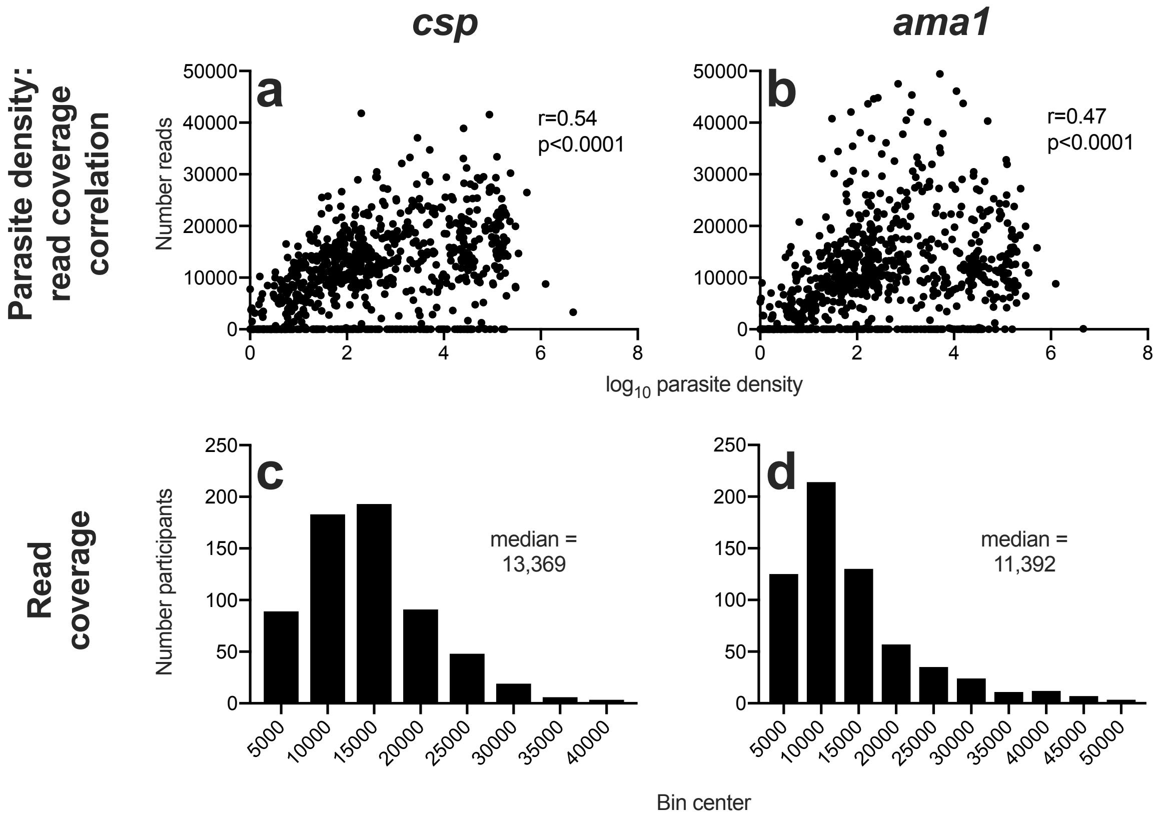

**Supplementary Fig. 2** Parasite density is well correlated with read coverage. **a,b** $Log_{10}$ parasite density for each sample is well correlated (Spearman rank test) with read coverage at *csp* (**a**) and *ama1* (**b**) loci. **c,d** Read coverage histograms for *csp* (**c**) and *ama1* (**d**) amplicons, indicating a median read coverage of 13,369 for the *csp* locus and 11,392 for the *ama1* locus.

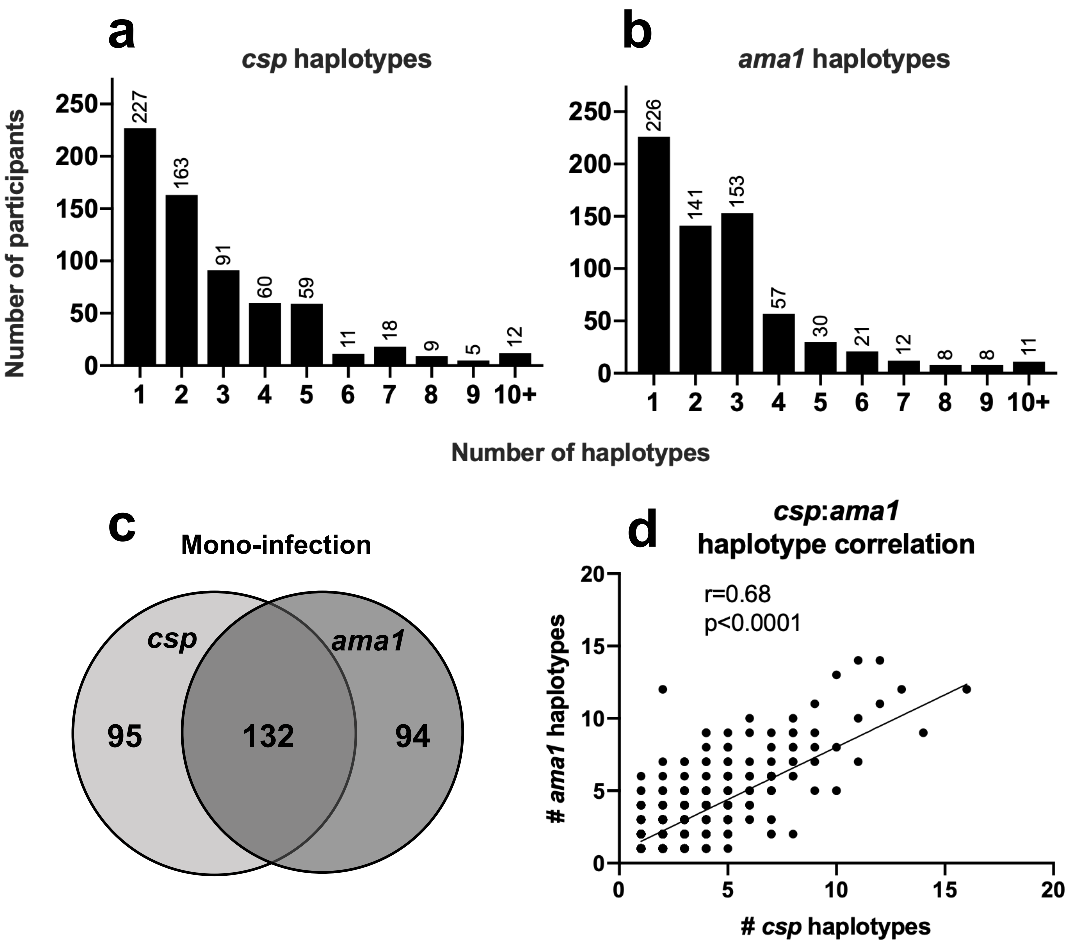

**Supplementary Fig. 3** Multigenomic infections detected in majority of study participants. **a,b** Histogram denoting the number of *csp* (a) and *ama1* (a) haplotypes identified in study participants. (C) Majority of samples with apparent *csp* mono-infection also have *ama1* mono-infection and vice versa. (D) Number of haplotypes identified at *csp* and *ama1* loci are well correlated.

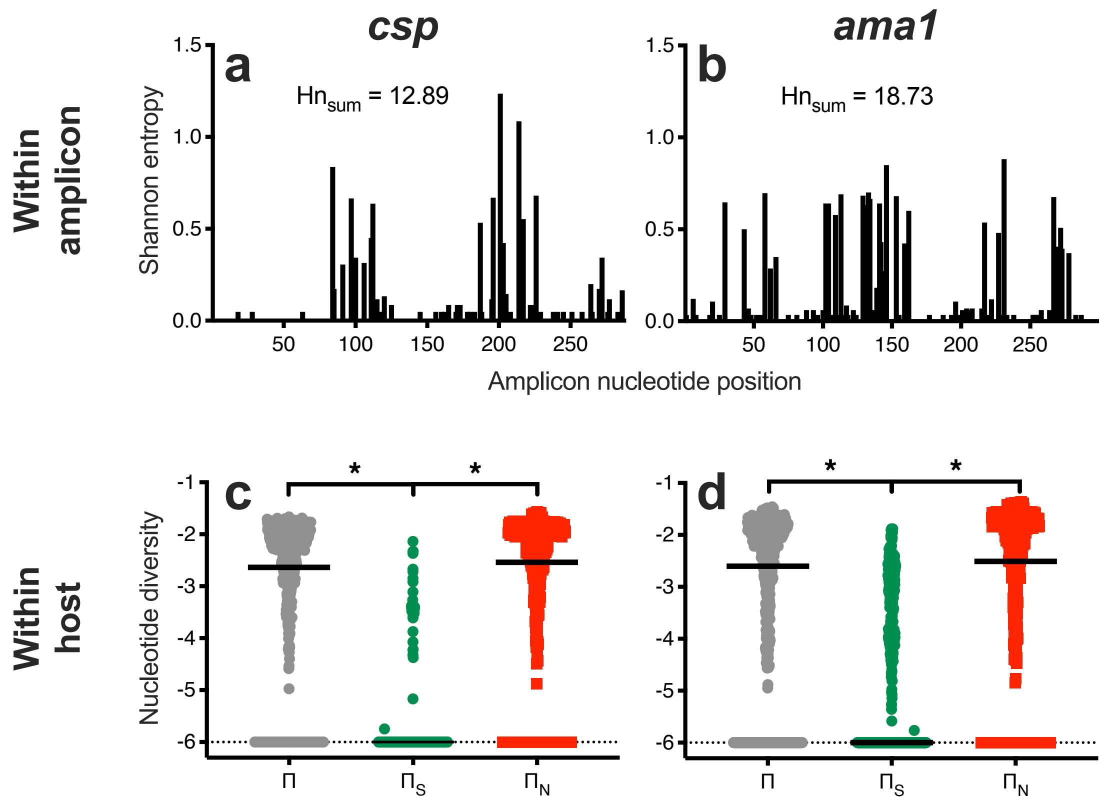

**Supplementary Fig. 4** Nucleotide diversity within amplicons and within hosts. **a,b** Shannon entropy scores at each position of the *csp* (**a**) and *ama1* (**b**) amplicons indicate that sequence diversity is restricted to 3 discrete regions within the csp amplicon though more evenly distributed in the *ama1* amplicon. Furthermore, the *ama1* amplicon has enhanced diversity overall with a total entropy ($Hn_{sum}$) of 18.73 compared with 12.89 for *csp.* **c,d** Intrahost nucletotide diversity ($\Pi$) is predominantly nonsynonymous ($\Pi_N$ ) rather than synonymous ($\Pi_S$) for both *csp* (**c**) and *ama1* (**d**) amplicons. * p<0.001, Friedman test + posthoc Wilcoxon Signed-Rank test.

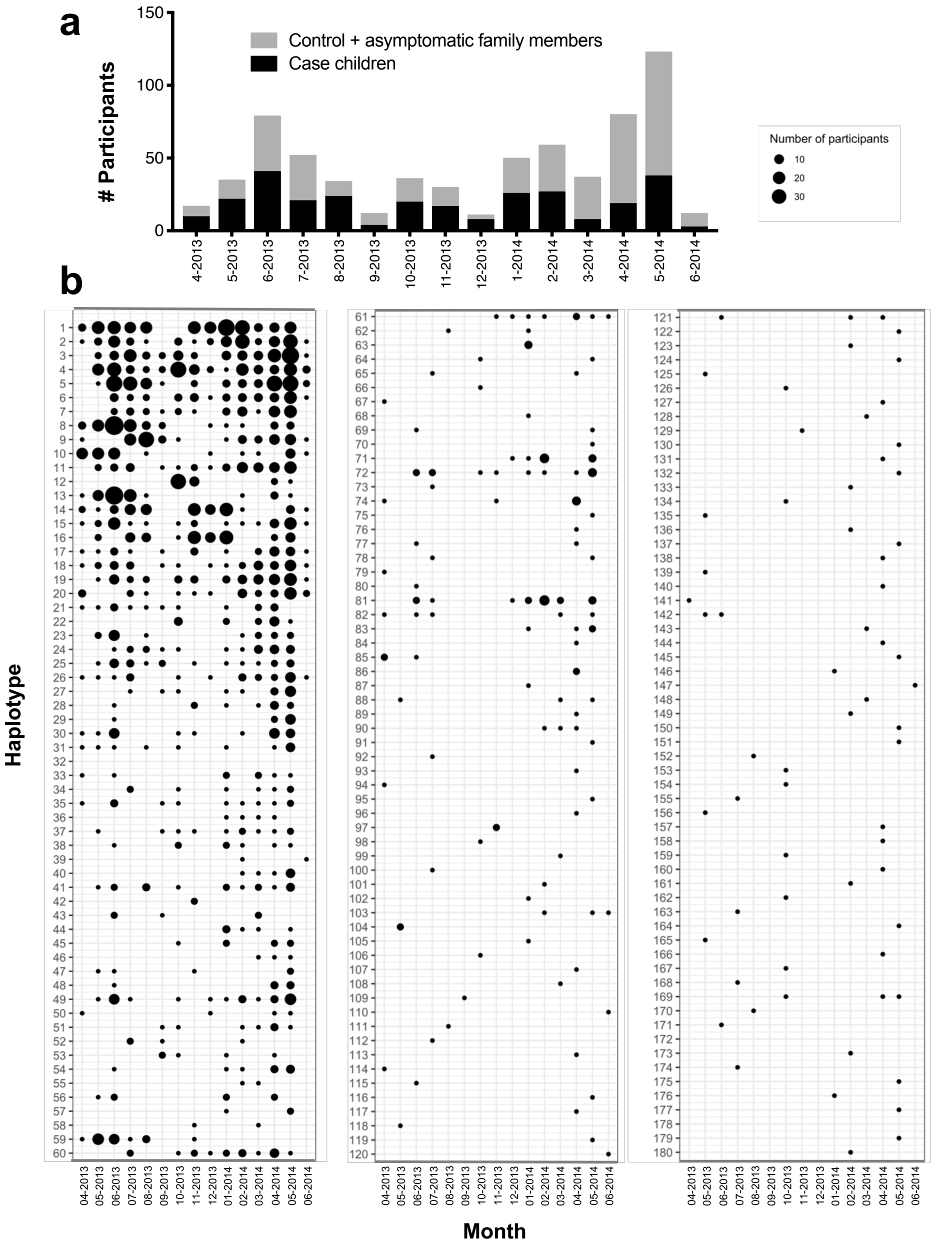

**Supplementary Fig. 5** *ama1* unique haplotype prevalence by month. **a** Total number of study participants with *ama1* haplotypes by month. Black denotes case children, and gray indicates both control and case household members. **b** Monthly prevalence of 180 unique *ama1* haplotypes, sorted by overall prevalence. Size of circle indicates number of study participants sharing a particular haplotype in a given month.

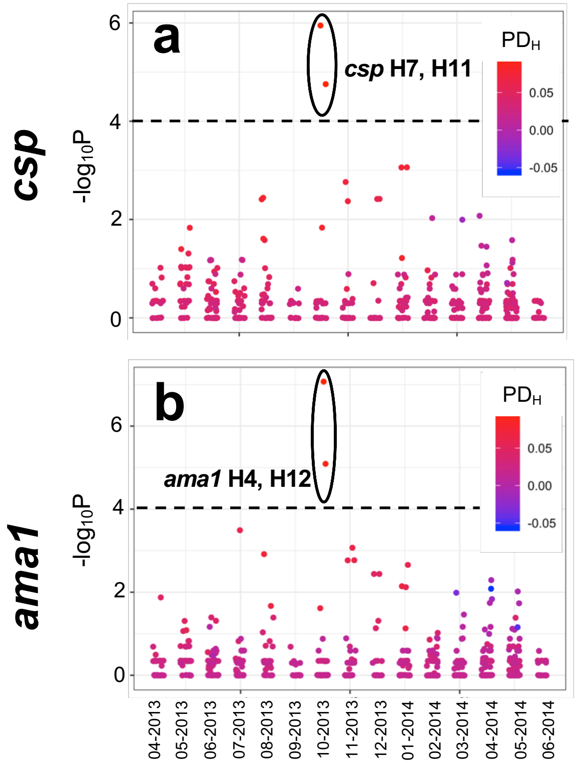

**Supplementary Fig. 6** No consistent haplotype bias by age **a,b** The haplotype prevalence difference ($PD_H$) between young children (≤ 5y; n=296 for *csp* and 300 for *ama1*) and older children/adults (>5y; n=357 for *csp* and 365 for *ama1*) during each month was calculated for *csp* (**a**) and *ama1* (**b**). Color indicates $PD_H$, with red identifying haplotypes more common in ≤ 5y population and blue more common in >5y population. The y-axis indicates the Fisher's Exact test $-\log_{10}$(p-value) for haplotype prevalence in ≤ 5y vs. >5y groups, while the dotted line denotes the Bonferroni-corrected threshold for statistical significance.

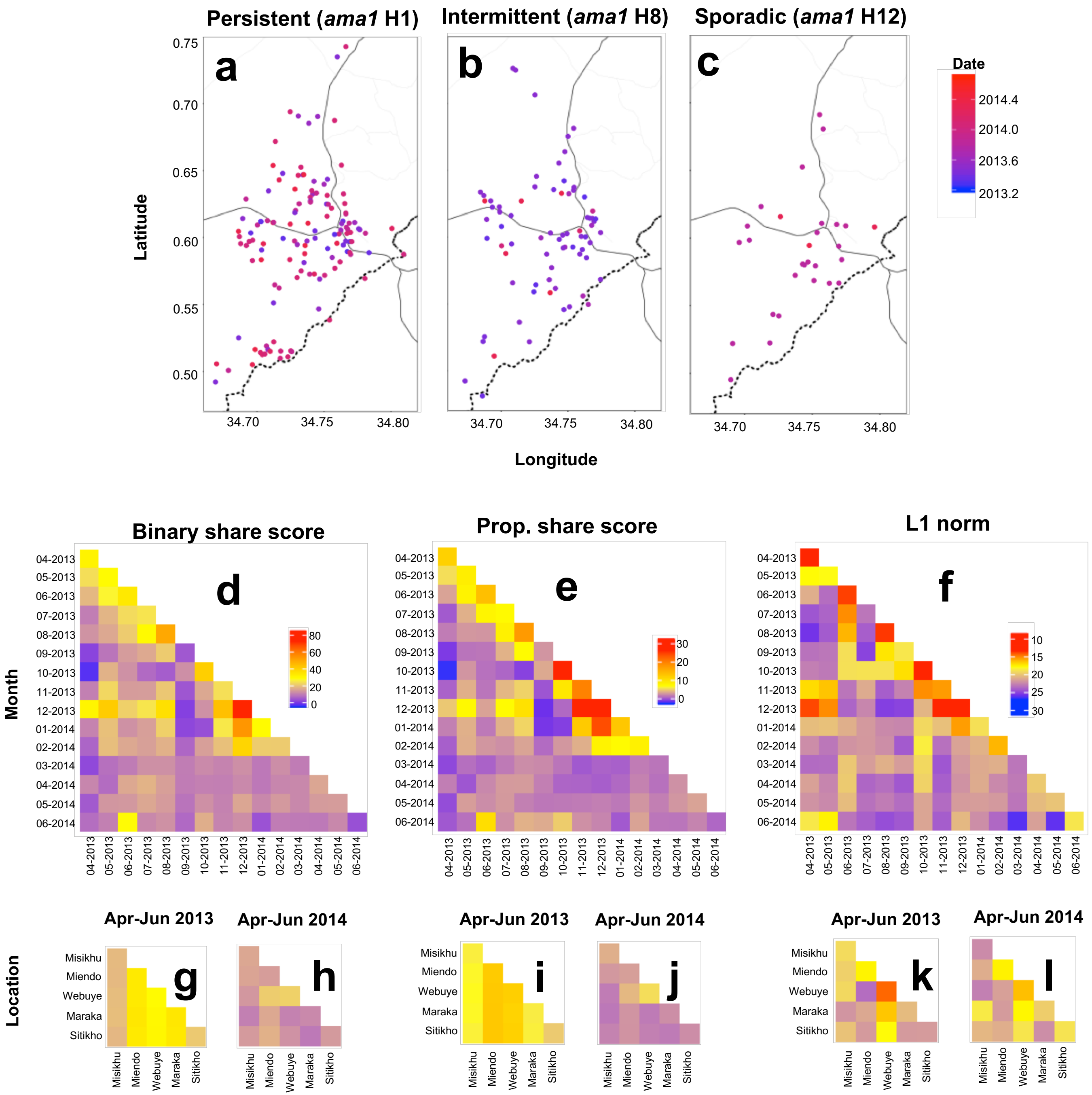

Persistent (ama1 H1)
Intermittent (ama1 H8)
Sporadic (ama1 H12)
a
b
c
Latitude
Longitude
Date
2014.4
2014.0
2013.6
2013.2
Binary share score
Prop. share score
L1 norm
d
e
f
Month
Apr-Jun 2013
Apr-Jun 2014
g
h
i
j
k
l
Location
Misikhu
Miendo
Webuye
Maraka
Sitikho

**Supplementary Fig. 7.** Genetic similarity of *ama1* haplotypes is structured by time more than space. **a-c** Location of study participants with ‘persistent’ haplotype *ama1* H1 (**a**), ‘intermittent’ haplotype *ama1* H8 (**b**), and ‘sporadic’ haplotype *ama1* H12 (**c**). Blue color indicates the beginning (April 2013) and red the end (June 2014) of the study period, with the date denoting fractional years in decimal notation. **d-f** Temporal comparison heat maps of mean binary haplotype sharing (**d**), proportional haplotype sharing (**e**), and L1 norm genetic distance (**f**) calculated between months of study enrollment. **g-l** Spatial comparison heat maps of binary haplotype sharing (**g,h**), proportional haplotype sharing (**i,j**), and L1 norm genetic distance (**k,l**) calculated for a distinct temporal window (**g,i,k**: April-June 2013; **h,j,l**: April-June 2014) for the 5 most represented administrative locations (see map in Figure S1), which are arranged geographically from north to south. For **d-l**, blue denotes the minimum for each genetic similarity index, red the maximum, and yellow the midpoint.

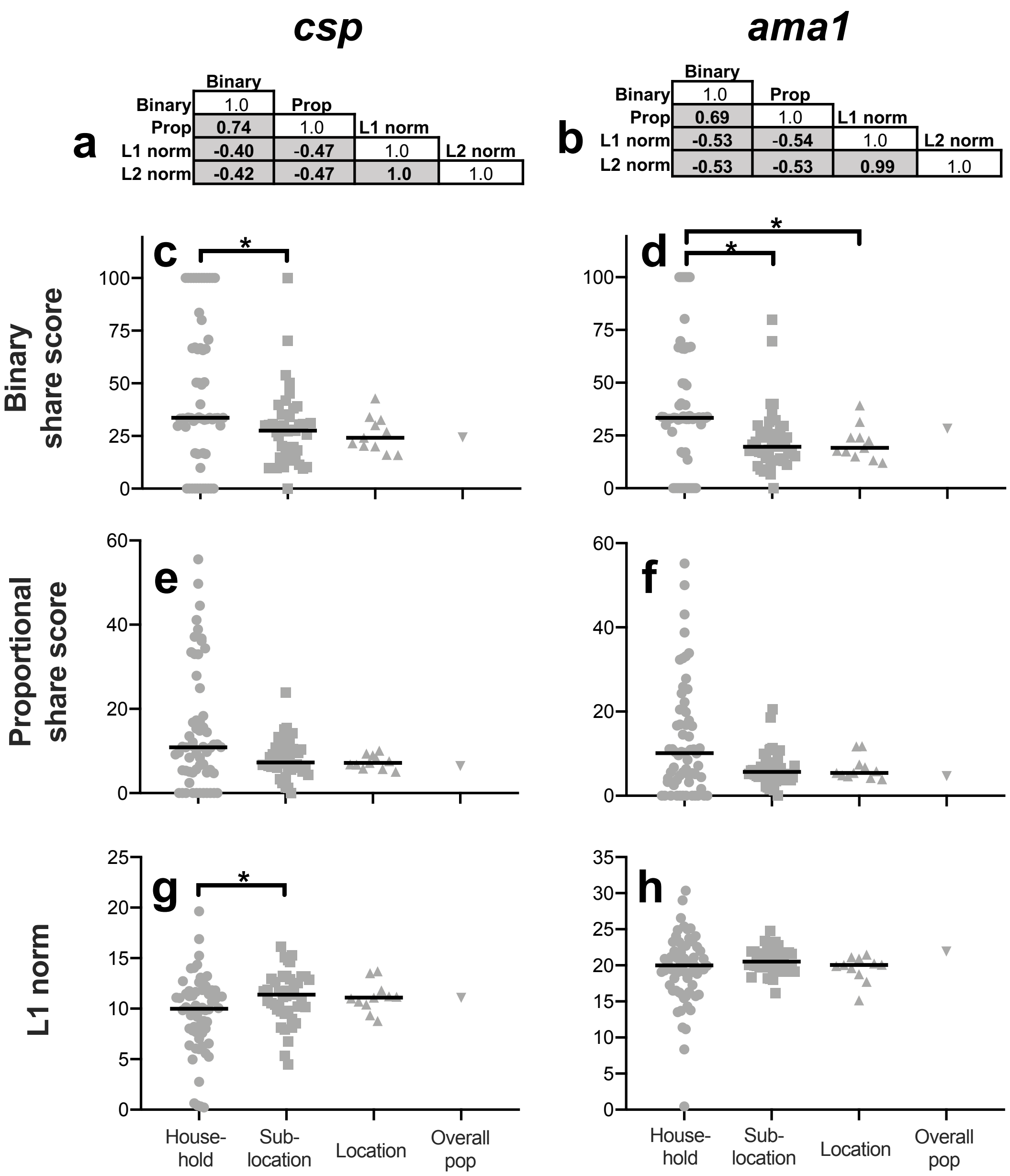

**Supplementary Fig. 8** Binary and proportional share scores are directly correlated, and inversely with L1/L2 norm metrics. **a,b** Correlation matrix (Spearman rank test) for binary share score, proportional share score (prop), L1 norm, and L2 norm calculated for *csp* (**a**) and *ama1* (**b**) loci. **c-h** Mean *csp* /*ama1* binary share score (**c,d**), proportional share score (**e,f**), and L1 norm (**g,h**) calculated for combination of individuals comprising each distinct household with 3+ members, sublocation, location, and for the overall population. *p<0.05, Mann-Whitney U test.

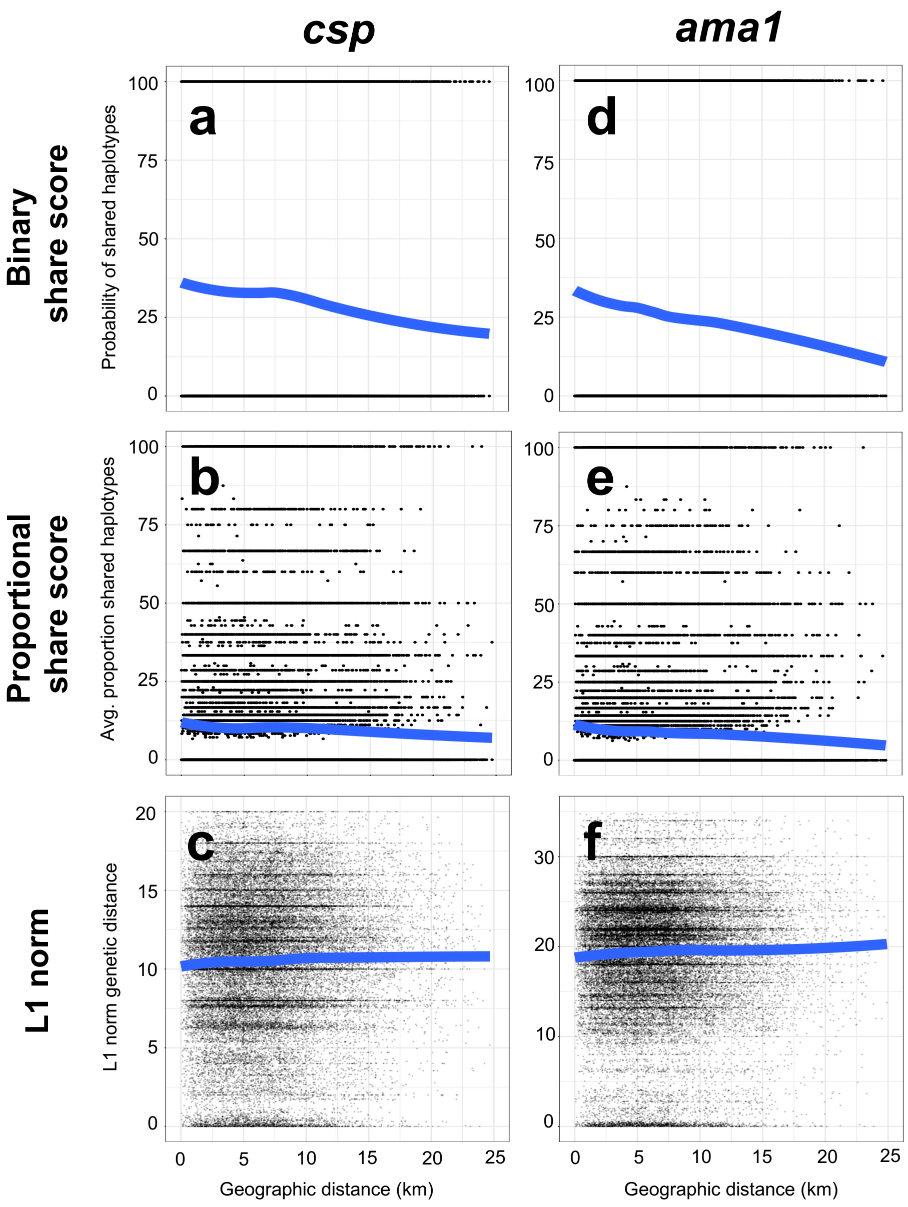

**Supplementary Fig. 9** No clear enhanced genetic similarity of infections among geographically-proximal symptomatic children. Genetic similarity metrics were computed for all possible pairings of case children for *csp* (n=273) and *ama1* (n=288) haplotypes. **a-c** *csp* haplotype binary sharing (**a**), proportional sharing (**b**) and L1 norm (**c**) metrics for all CC pairwise comparisons is plotted against geographic distance for CC. **d-f** *ama1* haplotype binary sharing (**d**), proportional sharing (**e**) and L1 norm (**f**) metrics for all CC pairings is plotted against geographic distance for CC. Blue lines indicate the locally-estimated scatterplot smoothing (LOESS) regression fit of data.

**Supplementary Table 1** Statistical comparison of study participants successfully/unsuccessfully assigned CSP and AMA haplotypes

| | | **Successful haplotype assignment** | **Unsuccessful haplotype assignment** | **P-value** | **Test** |
|---|---|---|---|---|---|
| | Median $log_{10}$ PD (range) | 2.36 (-0.51 – 6.67) | 0.71 (-0.86 – 5.85) | **p<0.001** | Mann-Whitney U Test |
| ***csp*** | Median age (range) | 6 (0.08 – 82) | 6 (0.08 – 73.5) | p=0.70 | Mann-Whitney U Test |
| | Case child percentage | 44.60% | 31.20% | **p<0.001** | Fisher's Exact Test |
| | Median $log_{10}$ PD (range) | 2.31 (-0.86 – 6.67) | 0.74 (-0.66 – 5.48) | **p<0.001** | Mann-Whitney U Test |
| ***ama1*** | Median age (range) | 6 (0.08 – 82) | 6 (0.08 – 73.5) | p=0.75 | Mann-Whitney U Test |
| | Case child percentage | 43.20% | 31.20% | **p<0.001** | Fisher's Exact Test |

**Supplementary Table 2** Comparison of genetic similarity metrics within-month

| | | **Binary sharing** | **Proportional sharing** | **L1-norm** |
|---|---|---|---|---|
| | Within-month median (n=15) | 36.17 | 13.61 | 8.51 |
| ***csp*** | Different month median (n=105) | 23.28 | 5.91 | 11.36 |
| | P-value (Mann-Whitney U test) | **0.029** | <**0.001** | **0.013** |
| | Within-month median (n=15) | 26.63 | 9.60 | 18.07 |
| ***ama1*** | Different month median (n=105) | 15.80 | 3.82 | 21.68 |
| | P-value (Mann-Whitney U test) | **0.004** | **<0.001** | **<0.001** |

**Supplementary Table 3** Comparison of genetic similarity metrics within-location

| | | Binary sharing | | Proportional sharing | | L1-norm | |
|---|---|---|---|---|---|---|---|
| | | Apr-Jun 2013 | Apr-Jun 2014 | Apr-Jun 2013 | Apr-Jun 2014 | Apr-Jun 2013 | Apr-Jun 2014 |
| ***csp*** | Within-location median (n=5) | 50.34 | 26.43 | 7.64 | 13.94 | 7.47 | 7.87 |
| | Different location median (n=10) | 53.69 | 18.34 | 4.12 | 15.11 | 8.96 | 8.96 |
| | P-value (Mann-Whitney U) | 0.594 | 0.076 | **0.002** | 0.371 | 0.240 | 0.165 |
| ***ama1*** | Within-location median (n=5) | 30.52 | 17.45 | 7.43 | 4.70 | 19.23 | 18.95 |
| | Different location median (n=10) | 31.57 | 16.87 | 9.95 | 3.69 | 20.52 | 20.79 |
| | P-value (Mann-Whitney U) | 1.000 | 0.514 | 0.207 | 0.075 | 0.207 | 0.254 |
